# Pathway-dependent regulation of sleep dynamics in a network model of the sleep-wake cycle

**DOI:** 10.1101/705822

**Authors:** Charlotte Héricé, Shuzo Sakata

## Abstract

Sleep is a fundamental homeostatic process within the animal kingdom. Although various brain areas and cell types are involved in the regulation of the sleep-wake cycle, it is still unclear how different pathways between neural populations contribute to its regulation. Here we address this issue by investigating the behavior of a simplified network model upon synaptic weight manipulations. Our model consists of three neural populations connected by excitatory and inhibitory synapses. Activity in each population is described by a firing-rate model, which determines the state of the network. Namely wakefulness, rapid eye movement (REM) sleep or non-REM (NREM) sleep. By systematically manipulating the synaptic weight of every pathway, we show that even this simplified model exhibits non-trivial behaviors: for example, the wake-promoting population contributes not just to the induction and maintenance of wakefulness, but also to sleep induction. Although a recurrent excitatory connection of the REM-promoting population is essential for REM sleep genesis, this recurrent connection does not necessarily contribute to the maintenance of REM sleep. The duration of NREM sleep can be shortened or extended by changes in the synaptic strength of the pathways from the NREM-promoting population. In some cases, there is an optimal range of synaptic strengths that affect a particular state, implying that the amount of manipulations, not just direction (i.e., activation or inactivation), needs to be taken into account. These results demonstrate pathway-dependent regulation of sleep dynamics and highlight the importance of systems-level quantitative approaches for sleep-wake regulatory circuits.

**Author Summary:** Sleep is essential and ubiquitous across animal species. Over the past half-century, various brain areas, cell types, neurotransmitters, and neuropeptides have been identified as part of a sleep-wake regulating circuitry in the brain. However, it is less explored how individual neural pathways contribute to the sleep-wake cycle. In the present study, we investigate the behavior of a computational model by altering the strength of connections between neuronal populations. This computational model is comprised of a simple network where three neuronal populations are connected together, and the activity of each population determines the current state of the model, that is, wakefulness, rapid-eye-movement (REM) sleep or non-REM (NREM) sleep. When we alter the connection strength of each pathway, we observe that the effect of such alterations on the sleep-wake cycle is highly pathway-dependent. Our results provide further insights into the mechanisms of sleep-wake regulation, and our computational approach can complement future biological experiments.

## Introduction

Global brain states vary dynamically on multiple timescales. Humans typically exhibit a daily cycle between three major behavioral states: wakefulness, REM sleep and NREM sleep. This daily cycle is regulated by a circadian rhythm and a homeostatic sleep pressure [1, 2]. These states alternate on a timescale of several hours called an ultradian rhythm [1, 3, 4]. Thus, complex interactions between homeostatic, circadian, and ultradian processes are involved in the sleep-wake cycle generation. However, it remains elusive how these states are regulated in the brain.

Over the past several decades, various cell types, neurotransmitters and neuropeptides have been identified as part of the sleep-wake regulating circuits within the brain [5–10]. Sleep- or wake-promoting neurons show state-dependent firing and contribute to the induction and/or maintenance of a particular state [5, 6, 8, 10–13]. To gain a better understanding of sleep-wake regulation, it is fundamental not just to identify and characterize each component of sleep-wake regulating circuits, but to also investigate how each pathway between neural populations contributes to state regulation.

Although controlling neural activity has provided mechanistic insights into sleep-wake regulation, their results are sometimes contradictory: for example, the role of pontine cholinergic neurons in REM sleep has been debated [14–17]. Even recent studies with opto- and chemogenetic approaches do not resolve this long-standing issue [17, 18]. Even though this discrepancy may be simply due to differences in animal models and experimental techniques, it is technically challenging to manipulate neurons or specific pathways precisely across different laboratories.

A computational approach may be a viable alternative for gaining insights into the mechanism of sleep-wake regulation. Since pioneering work in the 1970s and 80s [1, 3, 19], various computational models have been developed [3, 6, 20–25]: conceptually, a homeostatic sleep-dependent process and a circadian process play a key role in sleep regulation [1, 3]. Reciprocal excitatory-inhibitory connections [19, 21–23, 26] and mutual inhibitory interactions [8] can be recognized as key network motifs within sleep-wake regulating circuits. Although their dynamics have been explored [26–28], and those models can replicate sleep architecture of humans and animals [22] as well as state-dependent neural firing [25], few studies have investigated how the strength of synaptic connections between wake- and sleep-promoting populations contribute to sleep dynamics. As controlling neural activity at high spatiotemporal resolution *in vivo* becomes feasible, computational approaches can be considered as complementary approaches for investigating the role of specific neural pathways in sleep-wake regulation.

To this end, we utilize a simplified network model [26, 29] (**Figure 1**) and systematically manipulate the strength of every pathway. Because neurons within the sleep-wake regulating circuits typically project to a wide range of neural populations [6, 9, 30], their contributions to the sleep-wake cycle may also vary depending on the pathway. Therefore, we set out to test the hypothesis that the sleep-wake cycle is regulated in a pathway-dependent manner.

**Figure 1:**
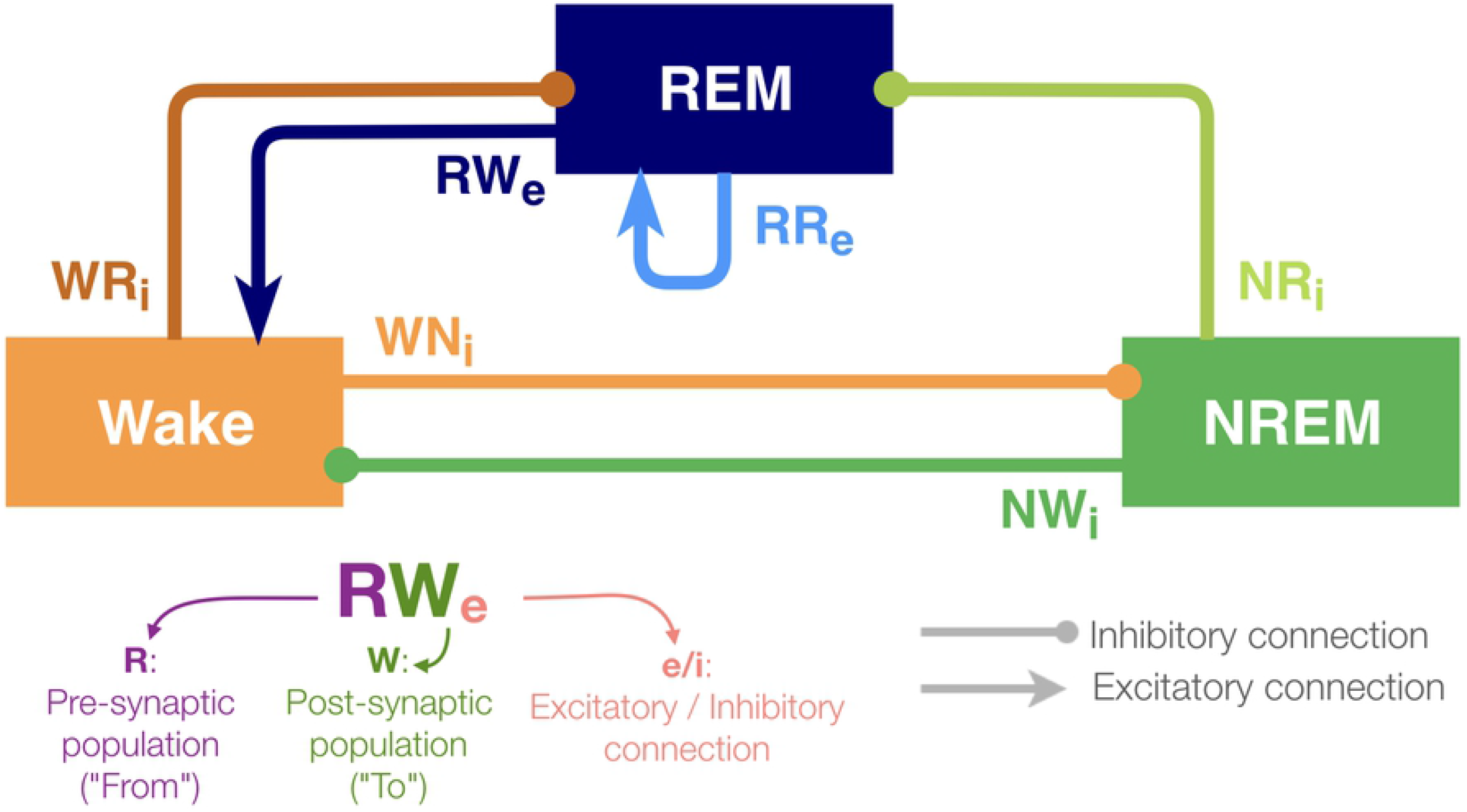
Architecture of the sleep regulatory network. Three neural populations are connected with excitatory and inhibitory synapses. Each neural population is named as the state they promote. The arrows and circles represent excitatory and inhibitory connections, respectively. The synapses are named with two uppercase and one lowercase letters: first letter of the pre-synaptic population (where the synapse is from), first letter of the post-synaptic population (where the synapse is going to) and “e” if it is excitatory or sign “i” if inhibitory.

Although the present model is highly abstract, it captures the following key features of sleep-wake regulating circuits: while the interaction between neuronal populations in the brainstem and the hypothalamus governs the sleep-wake regulation, some of the populations can be recognized as wake- or sleep-promoting [5–7, 9, 31]. To reflect the populations’ state-dependent firing, the model contains three neuronal populations (REM, NREM and Wake). The activity in these populations defines the state of the network (see Methods).

With respect to connectivity between these populations, Saper et al. proposed that the mutual inhibition between wake-promoting and sleep-promoting populations acts as a flip-flop switch for the regulation of transitioning between wakefulness and NREM sleep [8]. Hence, in this model, the outputs from the Wake-promoting and NREM-promoting populations are considered as inhibitory. Because pontine REM-active cholinergic neurons provide excitatory connections to the sublaterodorsal nucleus (SLD), a key component of REM sleep-regulating circuits [32], the REM-promoting model population has a recurrent excitatory connection. Glutamatergic neurons project rostrally to several wake-promoting nuclei, such as the intralaminar nuclei of the thalamus and basal forebrain, and the REM population also provides excitatory outputs onto the Wake population [32, 33]. In addition, because recent studies have shown that GABAergic inputs play a role in REM sleep induction [34], the REM-promoting population also receives inhibitory inputs from both the wake-promoting and NREM-promoting populations in this model. Based on this simplified model, we report that the effects of synaptic weight alterations on sleep architecture are highly pathway-dependent. We also discuss implications for future biological experiments.

## Results

We utilized the network architecture as reported in previous studies [26, 29]. As shown in **Figure 1**, this model contained three neuronal populations (labeled REM, NREM and Wake). The activity of these populations was characterized by differential equations describing the population firing rates which defined the state of the network (see Methods). These equations have been proved to be components of suitable sleep/wake regulatory computational models in previous studies [22, 23, 26, 27, 35]. The pathways from one population to the other were either excitatory or inhibitory. The concentrations of excitatory and inhibitory neurotransmitters were directly related to the population firing rates of the neural populations and a homeostatic sleep drive. Each population was perturbated by random Gaussian noise (**Supplementary Figure 1**).

### Sleep architecture under initial conditions

Before manipulating synaptic weights across pathways, we confirmed the sleep-wake cycle in our model (**Figure 2**). The initial parameter setting in our model was the same as that in previous reports [26, 29] (**Supplementary Table 1**). As shown in **Figure 2A**, this network always started with wakefulness where activity in the Wake-promoting population was high. As the homeostatic force gradually built up, the Wake-promoting population dropped its activity and the network entered NREM sleep. During sleep, the homeostatic force gradually decreased while alternations between NREM sleep and REM sleep appeared before the network exhibited wakefulness again. As expected, the concentration of neurotransmitters was well correlated with the firing rate of neural populations.

**Figure 2:**
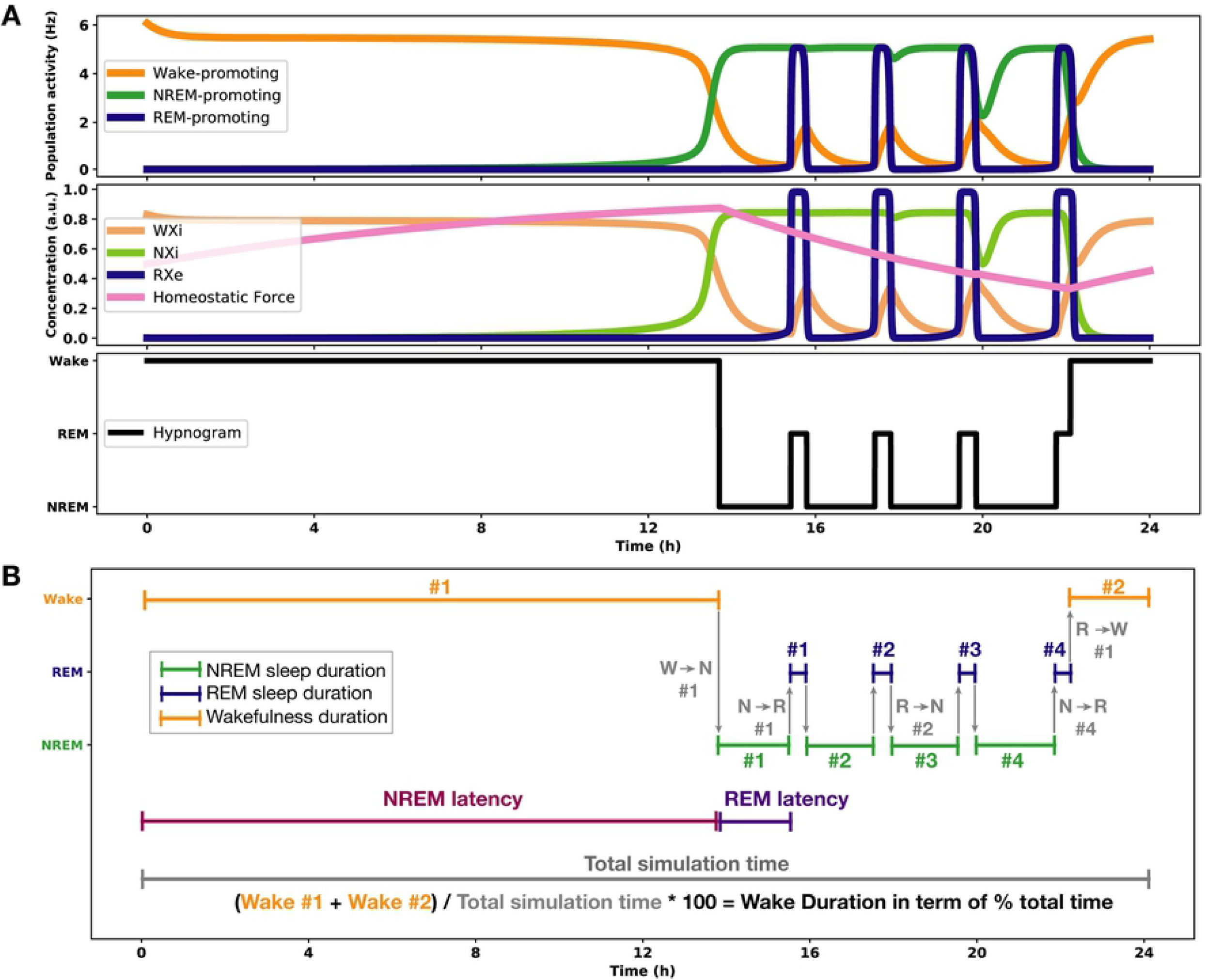
An example of the sleep-wake cycle generated by the network with the initial parameters. (**A**) top, the firing rate of each population as a function of time. Middle, the concentration of the neurotransmitters and the homeostatic force. Bottom, a hypnogram which was determined based on firing rates of the three neural populations. (**B**) Quantities of sleep architecture which were measured in this study The “#” stands for the episode or transition number in the order of appearance, i.e. W➔N #1 means the first transition from wakefulness to NREM.

In the following sections, to assess the effect of synaptic weight alterations on sleep architecture, we measured the following quantities (**Figure 2B**), all of which are measurable experimentally:

- the total duration of each state (**Figure 3 and Supplementary Figure 2**)
- the percentage of the time spent in these states (**Figures 4A, 5A, 6A**)
- the number of episodes (**Figures 4B, 5B, 6B**),
- the number of transitions between states (**Figures 4C, 5C, 6C**), and
- the NREM and REM sleep latencies (**Figure 7**).

In the following sections, we describe how synaptic weight alterations affect sleep architecture in this network with respect to these measurements.

### Effects of synaptic weight alteration on total sleep-wake duration

To investigate pathway-dependent regulation of sleep, we systematically modified the synaptic weight across pathways: the modified weight span from 0 to 8 times while *g* was the initial condition. We performed 24-hr simulations (n = 8) in each condition.

To assess the overall sleep architecture, we measured the total duration of each state (**Figure 3**). While each neural population had two output pathways (**Figure 1**), the effect of weight alterations on sleep architecture was highly pathway-dependent: in the case of the outputs from the Wake population, although stronger weights in the Wake ➔ NREM pathway led to longer wakefulness (*F*_1,7_ = 911.4, *p* < 0.0001, one-way ANOVA), the Wake ➔ REM pathway showed an opposite trend (*F*_1,7_ = 88.7, *p* < 0.0001, one-way ANOVA). The Wake ➔ NREM pathway was necessary to induce Wake whereas the Wake ➔ REM pathway was necessary to induce sleep states.

**Figure 3:**
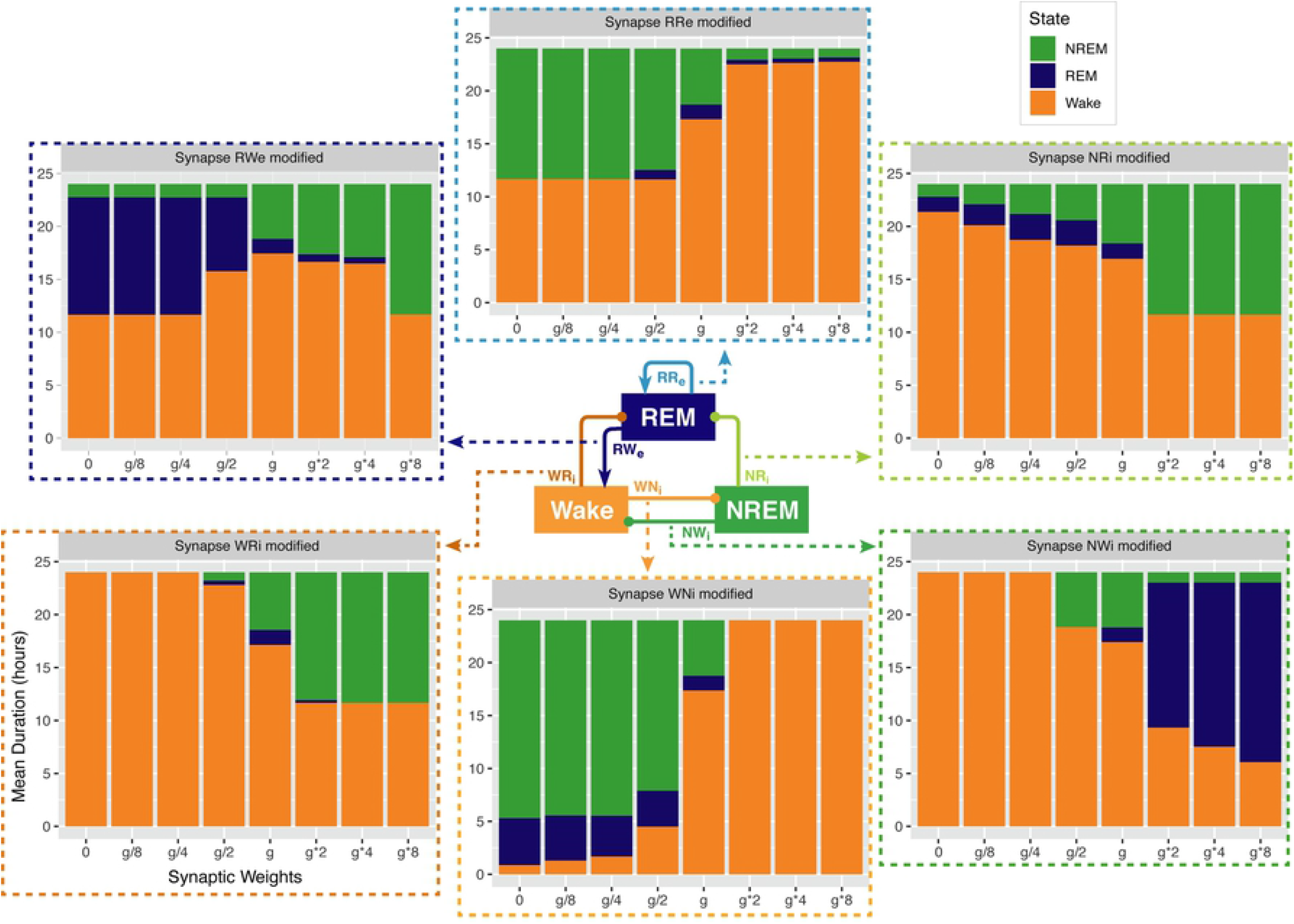
Total duration of each sleep state for different synaptic weights. Each bar graph represents the total duration of each state as a function of synaptic weights. The variable g represents the synaptic weight for the control condition. Each value is an average duration of each state from 8 simulations.

In the outputs from the NREM populations, stronger weights in the NREM ➔ REM connection led to longer NREM (*F*_1,7_ = 14985.8, *p* < 0.0001, one-way ANOVA) whereas stronger weights in the NREM ➔ Wake connection were associated with longer REM (*F*_1,7_ = 2290812, *p* < 0.0001, one-way ANOVA).

In the outputs from the REM population, to our surprise, strong recurrent excitatory connection shortened the duration of REM sleep (*F*_1,7_ = 189.2, *p* < 0.0001, one-way ANOVA). Rather, weaker weighting in the REM ➔ Wake connection promoted longer REM sleep (*F*_1,7_ = 94156.8, *p* < 0.0001, one-way ANOVA). Thus, the effects of synaptic weight alterations on overall sleep architecture were highly pathway-dependent. We also assessed how simultaneous alterations of two output pathways from each neural population affect sleep dynamics (**Supplementary Figure 3**). The outcomes deviated from those of individual pathway manipulations, suggesting pathway-dependent regulation in the sleep dynamics. In the next sections, we explore detailed sleep architecture of this model with varied synaptic weights.

### Alterations of REM population output pathways and overall sleep architecture

How does the output from the REM population contribute in the sleep architecture? To address this, we quantified the effect of synaptic weight alterations in the REM population outputs on the sleep architecture, with respect to the percentage of time spent in each state (**Figure 4A**), the number of episodes (**Figure 4B**), and the number of state transitions (**Figure 4C**).

**Figure 4:**
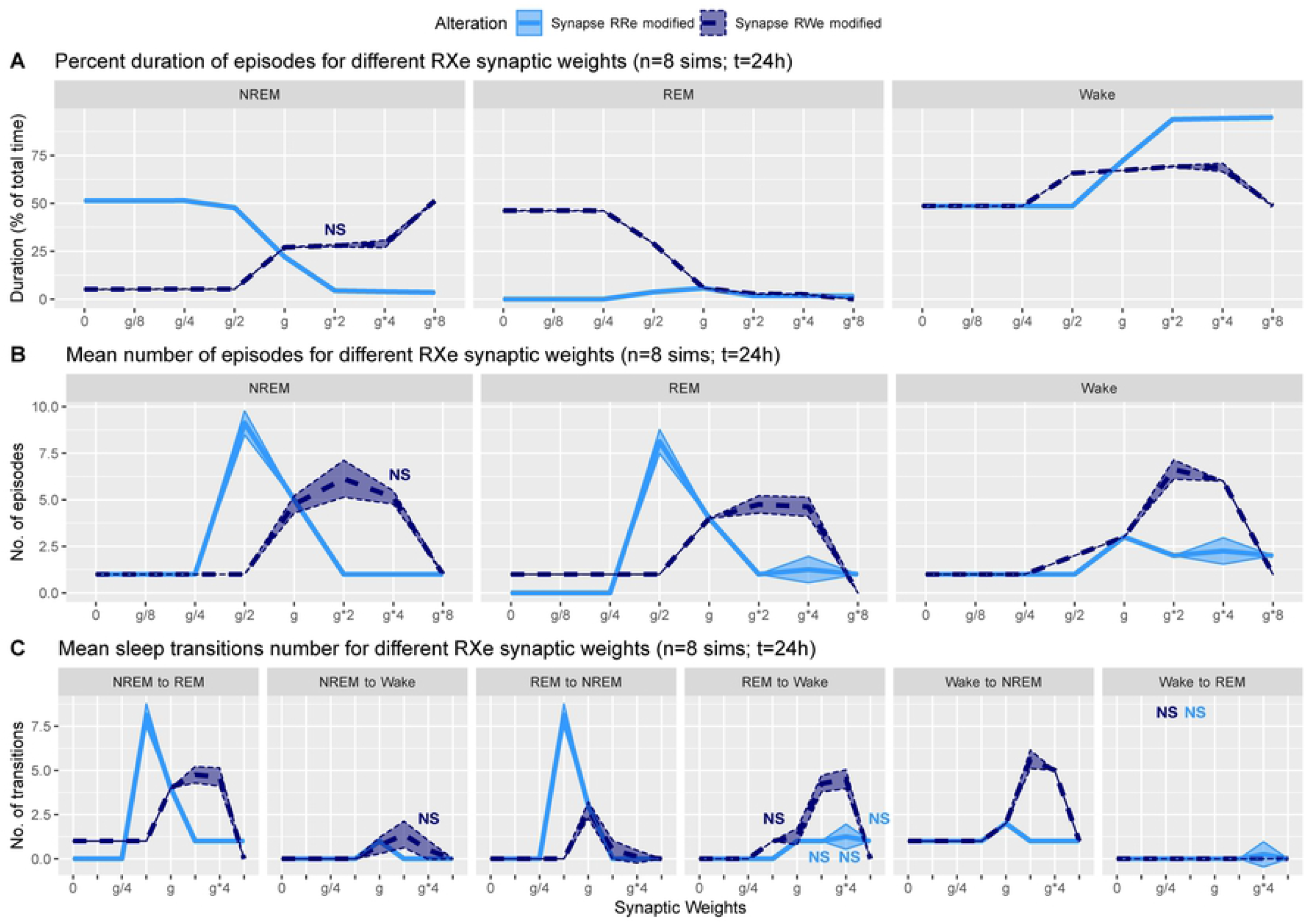
Effects of synaptic weight alterations of the REM population on sleep architecture. The percentage of time spent in each state (**A**), the number of episodes (**B**), and the number of state transitions (**C**) as a function of synaptic weights. Each profile was based on eight 24 hr simulations. Data presents mean ± s.e.m. Light blue, REM ➔ REM pathway; dark blue, REM ➔ Wake pathway. NS, non-significant (one-way ANOVA).

When we manipulated the synaptic weight in the REM ➔ REM excitatory recurrent pathway (light blue in **Figure 4**), the percentage of NREM sleep decreased as a function of the synaptic weight (*F*_1,7_ = 66736.6, *p* < 0.0001, one-way ANOVA) whereas the percentage of Wake increased (*F*_1,7_ = 67955.9, *p* < 0.0001, one-way ANOVA) (**Figure 4A**). We observed only small changes in the percentage of REM sleep. Although the percentage of time spent in NREM sleep was similar with weaker synaptic weights, the numbers of NREM and REM sleep episodes were exceptionally high only with *g*/2 (*F*_1,7_ = 1416.7 and *F*_1,7_ = 562.5 respectively, *p* < 0.0001 for both, one-way ANOVA) (**Figure 4B**). This was due to the increased number of transitions between NREM and REM sleep (*F*_1,7_ = 1255.9, *p* < 0.0001, one-way ANOVA) (**Figure 4C**). Because the number of Wake episodes was stable (**Figure 4B**), this particular synaptic weight induced fragmentation of sleep states.

When we manipulated the synaptic weight in the REM ➔ Wake excitatory pathway (dark blue in **Figure 4**), the percentage of REM sleep decreased as a function of the synaptic weight (*F*_1,7_ = 125513.9, *p* < 0.0001, one-way ANOVA) whereas the percentage of NREM sleep increased (*F*_1,7_ = 4543.5, *p* < 0.0001, one-way ANOVA) (**Figure 4A**). The weaker weight in the REM ➔ Wake pathway extended the duration of REM sleep (*F*_1,7_ = 189.2, *p* < 0.0001, one-way ANOVA). Although the time spent in REM sleep decreased with *g*\*2 and *g*\*4 (*F*_1,7_ = 9771.8, *p* < 0.0001, one-way ANOVA with post-hoc Tukey HSD test), the number of REM episodes (*F*_1,7_ = 497.1, *p* < 0.0001, one-way ANOVA) (**Figure 4B**) and transitions (**Figure 4C**) increased. Hence stronger REM ➔ Wake pathway caused a fragmented sleep-wake cycle although *g*\*8 provided a different picture, suggesting an optimal range of synaptic strengths to induce the fragmentation of the sleep-wake cycle. Therefore, effects of alterations of REM population output pathways on sleep architecture were highly pathway-dependent.

### Alterations of NREM population output pathways and sleep architecture

What are the effects of variation in the outputs from the NREM population in the sleep architecture and genesis? Here, we also examined how alterations of the output strengths from the NREM population contributed to sleep/wake transition, with respect to the percentage of time spent in each state (**Figure 5A**), the number of episodes (**Figure 5B**), and the number of state transitions (**Figure 5C**).

**Figure 5:**
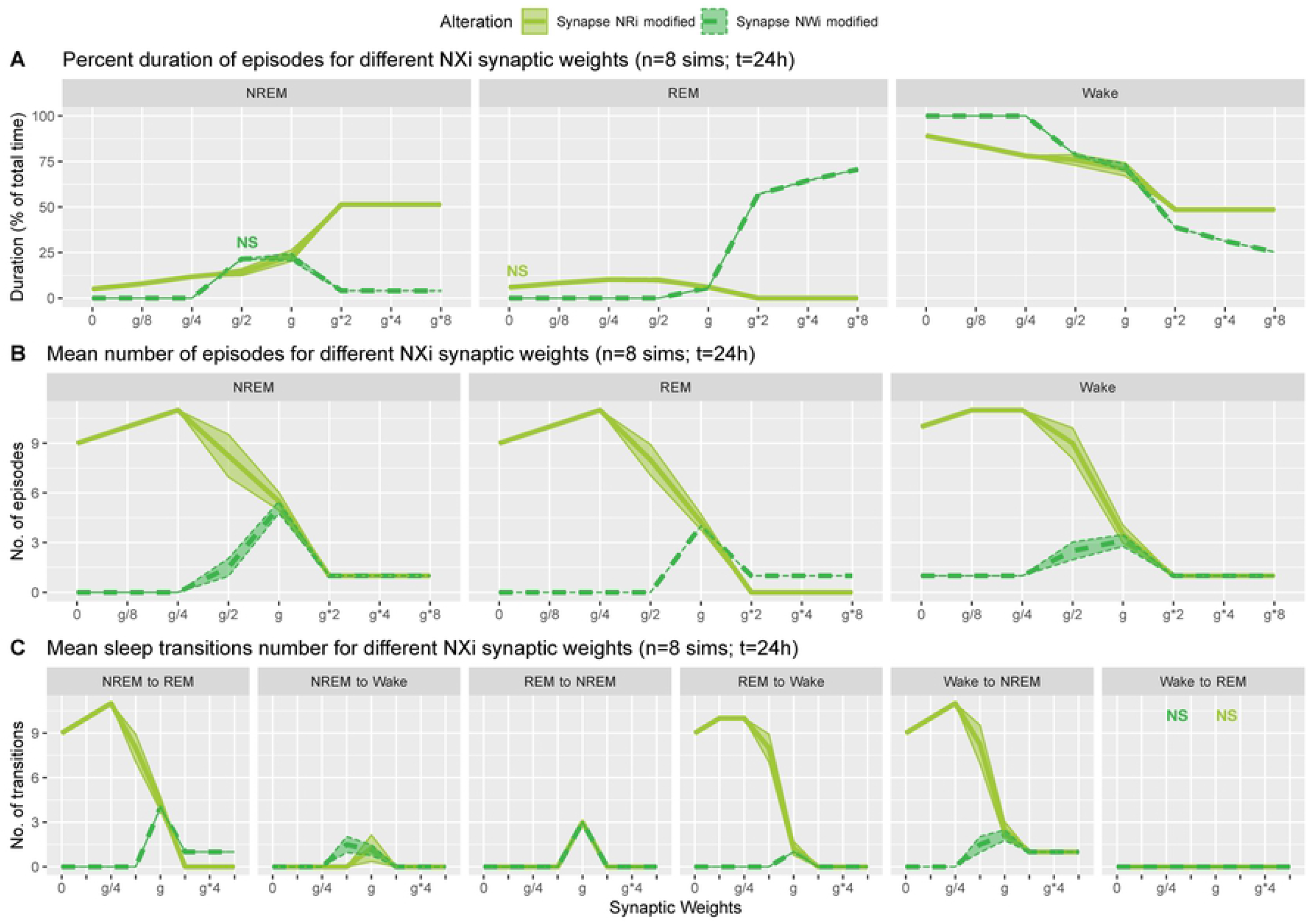
Effects of synaptic weight alterations of the NREM population on sleep architecture. The percentage of time spent in each state (**A**), the number of episodes (**B**), and the number of state transitions (**C**) as a function of synaptic weights. Each profile was based on eight 24 hr simulations. Data presents mean ± s.e.m. Light green, NREM ➔ REM pathway; green, NREM ➔ Wake pathway. NS, nonsignificant (one-way ANOVA).

Strengthening the NREM ➔ REM pathway (light green in **Figure 5**) increased the percentage of time spent in NREM (*F*_1,7_ = 2167.3, *p* < 0.0001, one-way ANOVA) and decreased that in REM (*F*_1,7_ = 867.9, *p* < 0.0001, one-way ANOVA) and Wake (*F*_1,7_ = 849.5, *p* < 0.0001, one-way ANOVA) (**Figure 5A**). This was associated with the reduction in state transitions (**Figures 5B and C**), meaning state stabilization. On the other hand, weakening the pathway increased the number of sleep episodes (*F*_1,7_ = 616.8, *p* < 0.0001, one-way ANOVA) and transitions (**Figures 5B and C**), meaning fragmentation.

Strengthening the NREM ➔ Wake pathway (green in **Figure 5**) increased the percentage of time spent in REM sleep (*F*_1,7_ = 165294.3, p < 0.0001, one-way ANOVA) and decreased that in NREM (*F*_1,7_ = 1543.9, p < 0.0001, one-way ANOVA) and Wake (*F*_1,7_ = 17388.3, p < 0.0001, one-way ANOVA) (**Figure 5A**). Weakening this pathway eliminated sleep episodes completely, meaning that this pathway is essential for sleep genesis.

### Alterations of Wake population output pathways and sleep architecture

We also examined how alterations of the output strengths from the Wake population contributed to sleep architecture, with respect to the percentage of time spent in each state (**Figure 6A**), the number of episodes (**Figure 6B**), and the number of state transitions (**Figure 6C**).

**Figure 6:**
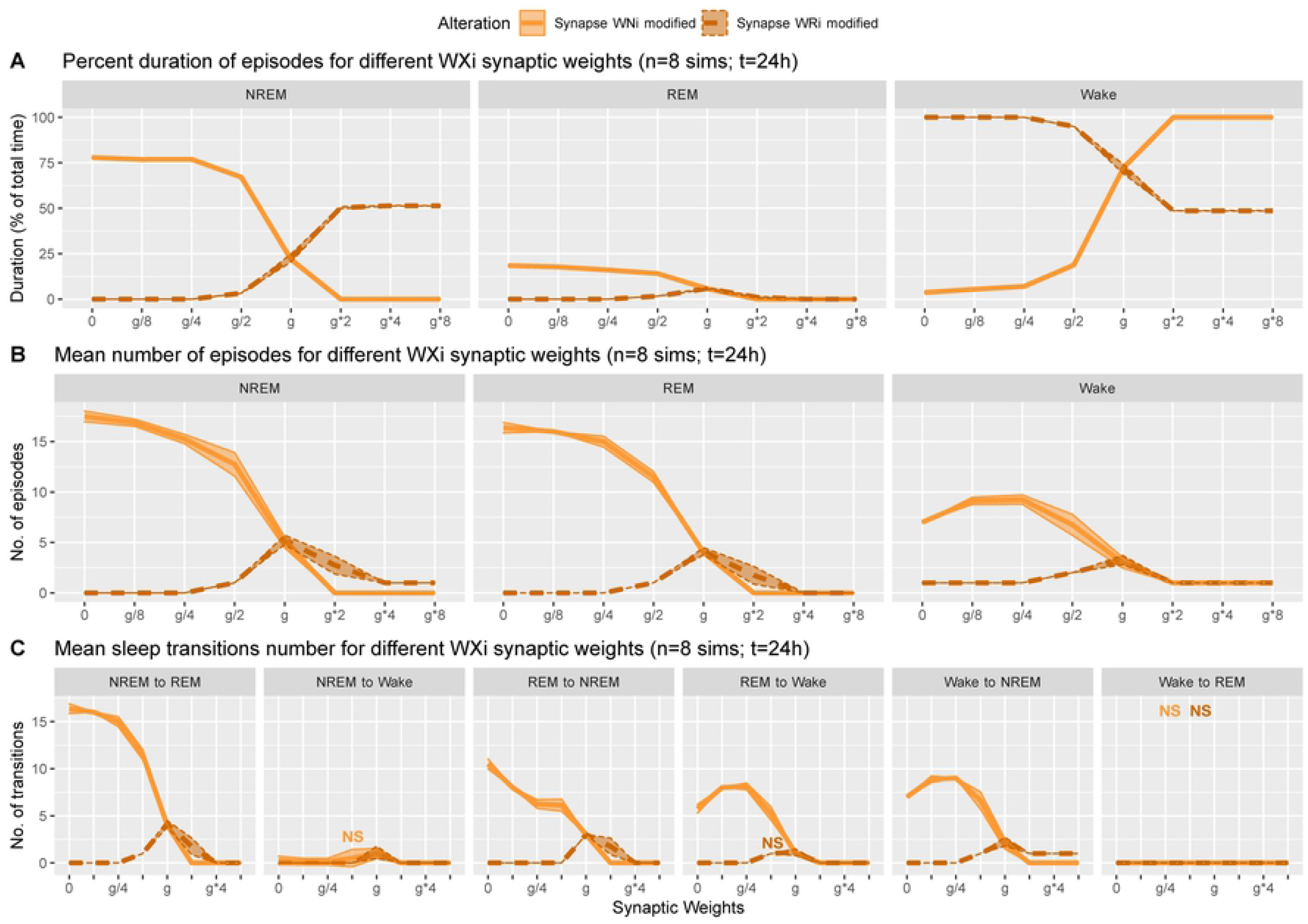
Effects of synaptic weight alterations of the Wake population on sleep architecture. The percentage of time spent in each state (**A**), the number of episodes (**B**), and the number of state transitions (**C**) as a function of synaptic weights. Each profile was based on eight 24 hr simulations. Data presents mean ± s.e.m. Orange, Wake ➔ NREM pathway; brown, Wake ➔ REM pathway. NS, non-significant (one-way ANOVA).

When we manipulated the synaptic weights in the Wake ➔ NREM inhibitory pathway (orange in **Figure 6**), the percentage of Wake increased as the synaptic weight increased (*F*_1,7_ = 99634.2, *p* < 0.0001, one-way ANOVA) (**Figure 6A**). On the other hand, as the synaptic weight decreased, the more the number of episodes increased across three states (*F*_1,7_ = 4391.6 for REM, *F*_1,7_ = 1788.8 for NREM, *F*_1,7_ = 504.9 for Wake, *p* < 0.0001 for all, one-way ANOVA) (**Figure 6B**), with longer sleep states (*F*_1,7_ = 20577.5, *p* < 0.0001, one-way ANOVA) (**Figure 6A**).

Contrary to these, when we increased the synaptic weight in the Wake ➔ REM pathway (brown in **Figure 6**), the percentage of Wake decreased (*F*_1,7_ = 5665.5, *p* < 0.0001, one-way ANOVA) (**Figure 6A**). There was an optimal range to increase the numbers of sleep episodes (*F*_1,7_ = 210.28, *p* < 0.0001, one-way ANOVA) (**Figure 6B**). Again, the effects of alterations of Wake population output pathways on sleep architecture were pathway-dependent.

### Effects of synaptic modifications on the sleep latency

We also measured the latency of NREM and REM (**Figures 2B and 7**): the former is the latency of the first NREM episode since the beginning of the simulation whereas the latter is the latency of the first REM episode since the onset of the first NREM episode.

**Figure 7:**
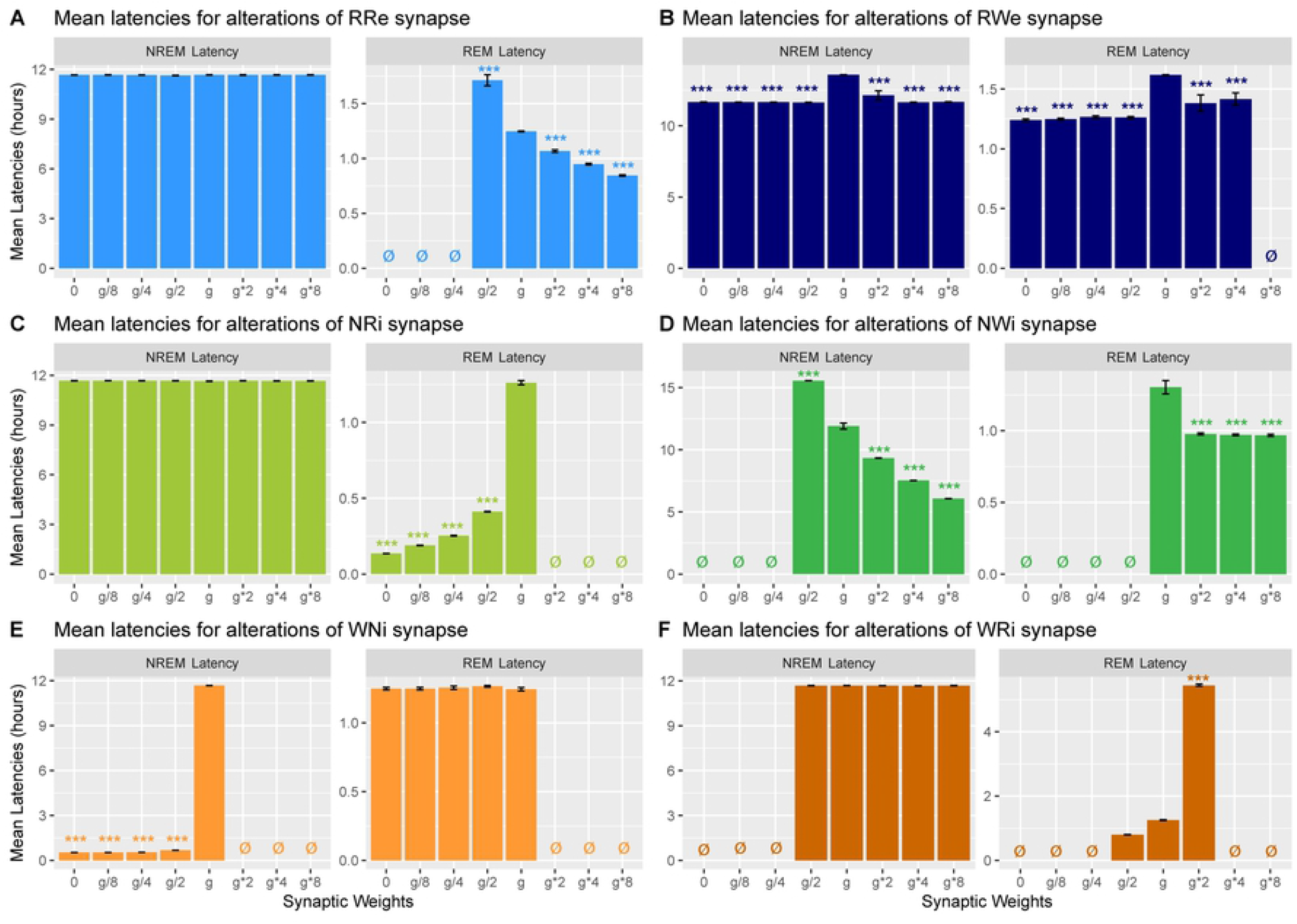
Effects of synaptic weight alterations on sleep latency. Bar graphs represent mean latency for NREM (left) and REM (right) as a function of synaptic weights in modifications of REM ➔ REM (A), REM ➔ Wake (B), NREM ➔ REM (C), NREM ➔ Wake (D), Wake ➔ NREM (E) and Wake ➔ REM pathways (F). Error bars, s.e.m.; ø, no occurrence of the state.

Strengthening the REM ➔ REM pathway decreased the REM latency (*F*_1,7_ = 202.5, *p* < 0.0001, one-way ANOVA) (**Figure 7A**) whereas strengthening the REM ➔ Wake pathway increased the REM latency only at g*4 (*F*_1,7_ = 17.3, *p* < 0.0001, one-way ANOVA with post-hoc Tukey HSD test) (**Figure 7B**). As expected, we did not observe any effect on the NREM latency by the manipulation of either pathway (**Figures 7A and B**). Thus, the output pathways from the REM population contributed only to the REM latency.

As expected, weakening the NREM ➔ REM inhibitory pathway decreased the REM latency (*F*_1,7_ = 4883, *p* < 0.0001, one-way ANOVA) whereas the NREM latency was not changed (**Figure 7C**). Strengthening the NREM ➔ Wake pathway decreased the NREM latency (*F*_1,7_ = 1199.2, *p* < 0.0001, one-way ANOVA) whereas the REM latency was also reduced and remained consistent across different weights (*F*_1,7_ = 47.3, *p* < 0.0001, one-way ANOVA) (**Figure 7D**). Thus, the output pathways from the NREM population exhibited complex contributions to the NREM and REM latencies depending on output pathways.

Finally, weakening the Wake ➔ NREM inhibitory pathway decreased the NREM latency (*F*_1,7_ = 303141.6, *p* < 0.0001, one-way ANOVA) whereas the REM latency was not affected as long as sleep was induced (**Figure 7E**). While strengthening the Wake ➔ REM inhibitory pathway did not affect the NREM latency, the REM latency increased at *g*2* (*F*_1,7_ = 16.3, *p* < 0.0001, one-way ANOVA with post-hoc Tukey HSD test). Thus, the output pathways from the Wake population contributed to the latency of sleep state which was directly influenced.

## Discussion

We report that the effect of synaptic weight manipulations on sleep architecture was highly pathway-dependent, based on the analysis of the simplified network model. Similar computational approaches can complement *in vivo* animal experiments to elucidate the circuit mechanisms of sleep-wake regulation.

In previous studies, the performances of network models have been explored [26–28] and these models can replicate sleep dynamics [22] as well as state-dependent neural firing [25]. Few studies have investigated how the strength of synaptic connections between wake- and sleep-promoting populations contribute to the sleep architectures. In this study, we systematically manipulated the strengths of neural pathways to investigate the effect on sleep architectures.

Our model exhibited several non-trivial behaviors. For example, the duration of REM sleep was prolonged by some intuitive manipulations that directly involve the REM-promoting populations (**Figures 3 and 4**) such as: 1) weakening the REM ➔ Wake excitatory pathway and 2) weakening the NREM ➔ REM inhibitory pathway. In these two cases, each manipulation directly opposes the transition from REM sleep to other states. Surprisingly, we also observed this prolongation of REM sleep when the REM population is not directly involved in the manipulations, for example: 3) weakening the Wake ➔ NREM inhibitory pathway, or 4) strengthening the NREM ➔ Wake inhibitory pathway.

Given the simple network architecture (**Figure 1**), it may not be surprising to see such effects: that is, the pathways which are not directly connected to the REM population can contribute to the duration of REM sleep. However, these observations have an important implication for designing *in vivo* experiments and interpreting the outcomes. It is possible that any manipulations can make distal impacts, resulting in unexpected state alternations. This effect is called “Diaschisis” or “shocked throughout”, describing the sudden loss of function in another portion of the brain through being linked with a distal, (directly) affected brain region [36, 37].

Similarly, the dependency on manipulated pathways also has important implications. For example, when strengthening an output pathway from the NREM population, opposite effects on the duration of NREM sleep can be seen depending on the manipulated pathway. Thus, in highly interconnected circuits like sleep-wake regulating circuits, effects of manipulations on the system’s behavior can be differed by changing the manipulated pathways. In this area, systematic quantitative approaches are essential.

Another intriguing observation from our results is that measuring the latency of sleep states provided relatively intuitive outcomes. For example, strengthening the REM ➔ REM recurrent excitatory pathway could reduce the REM latency without increasing the duration of REM sleep. Strengthening the NREM ➔ Wake inhibitory pathways also reduced the NREM latency. Thus, measuring the latency to change states may provide insights into the role of the manipulated pathway in sleep regulation.

One of the major limitations in the present study is that the network model did not fully capture biological sleep-wake regulation. Therefore, some of our observations do not necessarily predict the behavior of biological circuits. To address this, it would be important to extend the network size to reflect more biological observations [25]. However, more quantitative experimental data are required to create even more realistic networks [6].

Another limitation to the present work is that we manipulated the synaptic strength during the entire simulation period. In biological experiments, however, manipulations can be transient, such as in optogenetic experiments [17, 34, 38]. It would be interesting to manipulate synaptic weights transiently in the network model.

Relating to this point, it may be also interesting to reconsider the definition of the state in the model. In particular, if the activity of each neural population is manipulated, the current definition (see Methods) can not be adopted, because the activity of each population itself defines the state. One way to address this issue is by connecting the modeled sleep-wake regulating circuit to cortical circuits and muscle units, through which the state of the system can be defined based on the activity of the cortical circuits or muscle units as in biological experiments, where cortical electroencephalograms and electromyograms are used to define behavioral states.

In conclusion, utilizing a simple network model of the sleep-wake cycle, we found pathway-dependent effects of synaptic weight manipulations on sleep architecture. Given the fact that even the simple network model can provide complex behaviors, designing *in vivo* experiments and interpreting the outcomes to require careful considerations about the complexity of sleep-wake regulating circuits. A similar computational approach with a biologically realistic network could complement to make specific predictions for *in vivo* experiments.

## Methods

We implemented a computational model of the sleep/wake cycle containing three neuronal populations whose activity by several differential equations. Numerical simulations were computed with the Runge-Kutta integration method (4^th^ order), with a time step of 1ms and a simulation duration of 24h. For these simulations and a part of the data processing, we used the Python programming language (version 3.6.8). In order to run multiple simulations for all the conditions, we implemented a script Bash (version 3.2.57). The majority of the data processing, the plots were performed with R (version 3.5.1). All details about the tools and libraries used for this work are summarized in **Supplementary Table 2**. All codes are available at https://github.com/SakataLab/Sleep_Computational_Model_2019.

### Firing rate formalism

All three populations are promoting the sleep-wake cycle corresponding to their name and are associated with a specific neurotransmitter. The REM-promoting population releases the excitatory neurotransmitters RX_e_ whereas the NREM- and Wake-promoting populations release the inhibitory ones NX_i_ and WX_i_, respectively.

Firing rate *F_X_* of population *X* is described as follows:

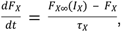

where *F*_*X*∞_ is a steady state firing rate function for each population (see below). *τ*_X_ is the membrane time constant of the population *X*. The synaptic input *I_X_* is a weighted sum of neurotransmitter concentrations released by the pre-synaptic populations *Y* and is described as follows:

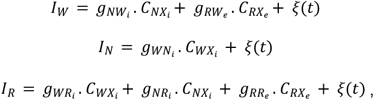

where *C_YXe/i_* represents the neurotransmitter concentration involved in the pathway from population *Y* to *X* with synaptic weight *g_YXe/i_*. The parameter *ξ*(*t*) provides a weak Gaussian noise (mean of 0.01 Hz and standard deviation of 0.005 Hz) to mimics the variability of the biological networks.

For each population, the steady state firing rate function *F*_*X*∞_ is modelled with the following equations:

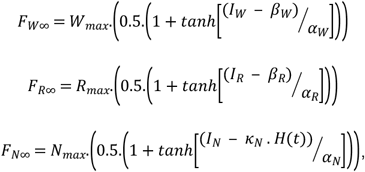

where W_max_, N_max_ and R_max_ are constant values to set the maximum firing rates of the populations. α and β are slope and threshold parameters of the hyperbolic tangent function for the population X, respectively. Because the NREM population is linked to the homeostatic sleep drive, the steady state firing function also depends on the homeostatic sleep drive variable *H*(*t*) (see below).

All parameter values are provided in **Supplementary Table S1**.

### Homeostatic sleep drive

In the model, the sleep-wake transition is driven by the homeostatic sleep drive *H*(*t*). This process can be described by the following equation:

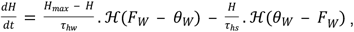

where 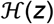 stands for the Heaviside function, which returns 0 if z < 0 and 1 if z ≥ 0. *θ_W_* is a constant to set the sleep drive threshold. *H_max_* is a constant value to set the maximum value for the homeostatic force. T_*hw*_ and T_*hs*_ are time constants of sleep drive built up during wakefulness and declined during sleep, respectively. Hence, the homeostatic force value increases during wakefulness due to a high activity of the wake-promoting population, and decreases during sleep when this activity is lower.

### Action of neurotransmitters

The neurotransmitter concentration *C_YX_*(*t*) from the populations Y to X is described as following:

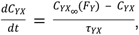

where *C*_*YX*∞_ is a saturating function to provide the steady state of the neurotransmitter release from the population Y to the population X as a function of F_Y_. This function is described as:

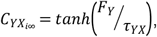

where T_*YX*_ is a time constant. The concentration of each neurotransmitter was normalized between 0 and 1 and is expressed in arbitrary unit (a.u.) [22].

### Alterations of synaptic weights in the network

To investigate pathway-dependent regulation of sleep architecture in the network model, we systematically altered the synaptic weight *g* in the network as shown in **Table 1**.

**Table 1:** Synaptic weights for the different alterations. Initials values can be found in the Supplementary Table S2 with the model parameters.

| Conditions | Eighth | Quarter | Half | Double | Quadruple | Octuple |
| --- | --- | --- | --- | --- | --- | --- |
| Symbols | $g/8$ | $g/4$ | $g/2$ | $g*2$ | $g*4$ | $g*8$ |
| $RR_e$ | 0.2 | 0.4 | 0.8 | 3.2 | 6.4 | 12.8 |
| $RW_e$ | 0.125 | 0.25 | 0.5 | 2.0 | 4.0 | 8 |
| $WN_i$ | -0.25 | -0.5 | -1.0 | -4.0 | -8.0 | -16.0 |
| $WR_i$ | -0.5 | -1.0 | -2.0 | -8.0 | -16.0 | -32.0 |
| $NR_i$ | -0.1625 | -0.325 | -0.65 | -2.6 | -5.2 | -10.4 |
| $NW_i$ | -0.21 | -0.42 | -0.84 | -3.36 | -6.72 | -13.44 |

We also simulated a lesion of each pathway by setting *g* to 0. For each condition, we run 8 simulations.

### Determination of sleep-wake states

The state of the network was determined according to Diniz Behn and Booth (2010): If firing rate of the Wake-promoting population is above 2 Hz, the state of the network is Wake. If not, the state is either NREM or REM sleep: if firing rate of the REM-promoting population is above 2 Hz, the state is REM sleep. If not, the state is NREM sleep.

### Statistical Analyses

All statistical analyses were performed using R scripts (version 3.5.1). Data are presented as the means (plain curves) ± s.e.m. (shaded curves). One-way analysis of variance (ANOVA) were used to analyse the synaptic weights alterations depending on the sleep state or transition. Following the ANOVA, Tukey *post-hoc* tests were performed for pairwise comparisons to the control conditions (no synaptic weights manipulations). *P*-values less than 0.05 were considered significant. If it is not the case, the sign “NS” was added on the graphs, otherwise there was a significant difference compared to the control condition.

## Author Contributions

SS and CH designed the project. CH developed the code, performed the simulations and analyzed the data. SS and CH wrote the manuscript.

## Acknowledgments

We thank Arno Onken, William Berg, Aimee Bias and Emma Mitchell for critical reading of a manuscript. This work was supported by BBSRC (BB/M00905X/1), Leverhulme Trust (RPG-2015-377) and Alzheimer’s Research UK (ARUK-PPG2017B-005) to SS.

## References

1. Borbely AA. A two process model of sleep regulation. Hum Neurobiol. 1982;1(3):195–204. Epub 1982/01/01. PubMed PMID: 7185792.

2. Achermann P, Borbely AA. Simulation of human sleep: ultradian dynamics of electroencephalographic slow-wave activity. J Biol Rhythms. 1990;5(2):141–57. Epub 1990/01/01. doi: 10.1177/074873049000500206. PubMed PMID: 2133124.

3. Archermann P, Borbely AA. Sleep Homeostasis and Models of Sleep Regulation. In: Kryger M, Roth T, Dement WC, editors. Principles and Practice of Sleep Medicine. 6th Edition ed. Philadelphia: Elsevier; 2017. p. 377–87.

4. Carskadon MA. Normal Human Sleep: An Overview. In: Kryger M, Roth T, Dement WC, editors. Principles and Practice of Sleep Medicine. 6th edition ed. Philadelphia: Elsevier; 2017. p. 15–24.

5. Brown RE, Basheer R, McKenna JT, Strecker RE, McCarley RW. Control of sleep and wakefulness. Physiol Rev. 2012;92(3):1087–187. doi: 10.1152/physrev.00032.2011. PubMed PMID: 22811426; PubMed Central PMCID: PMCPMC3621793.

6. Herice C, Patel AA, Sakata S. Circuit mechanisms and computational models of REM sleep. Neurosci Res. 2019;140:77–92. Epub 2018/08/18. doi: 10.1016/j.neures.2018.08.003. PubMed PMID: 30118737; PubMed Central PMCID: PMCPMC6403104.

7. Luppi PH, Clement O, Fort P. Paradoxical (REM) sleep genesis by the brainstem is under hypothalamic control. Curr Opin Neurobiol. 2013;23(5):786–92. doi: 10.1016/j.conb.2013.02.006. PubMed PMID: 23490549.

8. Saper CB, Chou TC, Scammell TE. The sleep switch: hypothalamic control of sleep and wakefulness. Trends Neurosci. 2001;24(12):726–31. Epub 2001/11/24. PubMed PMID: 11718878.

9. Scammell TE, Arrigoni E, Lipton JO. Neural Circuitry of Wakefulness and Sleep. Neuron. 2017;93(4):747–65. doi: 10.1016/j.neuron.2017.01.014. PubMed PMID: 28231463; PubMed Central PMCID: PMCPMC5325713.

10. Weber F, Dan Y. Circuit-based interrogation of sleep control. Nature. 2016;538(7623):51–9. doi: 10.1038/nature19773. PubMed PMID: 27708309.

11. Hobson JA, McCarley RW, Wyzinski PW. Sleep cycle oscillation: reciprocal discharge by two brainstem neuronal groups. Science. 1975;189(4196):55–8. PubMed PMID: 1094539.

12. Jouvet M. [Research on the neural structures and responsible mechanisms in different phases of physiological sleep]. Arch Ital Biol. 1962;100:125–206. PubMed PMID: 14452612.

13. McCarley RW, Hobson JA. Single neuron activity in cat gigantocellular tegmental field: selectivity of discharge in desynchronized sleep. Science. 1971;174(4015):1250–2. PubMed PMID: 5133450.

14. Grace KP. How useful is optogenetic activation in determining neuronal function within dynamic circuits? Proc Natl Acad Sci U S A. 2015;112(29):E3755. Epub 2015/07/01. doi: 10.1073/pnas.1506188112. PubMed PMID: 26124088; PubMed Central PMCID: PMCPMC4517253.

15. Grace KP, Horner RL. Evaluating the Evidence Surrounding Pontine Cholinergic Involvement in REM Sleep Generation. Front Neurol. 2015;6:190. Epub 2015/09/22. doi: 10.3389/fneur.2015.00190. PubMed PMID: 26388832; PubMed Central PMCID: PMCPMC4555043.

16. Grace KP, Vanstone LE, Horner RL. Endogenous cholinergic input to the pontine REM sleep generator is not required for REM sleep to occur. J Neurosci. 2014;34(43):14198–209. Epub 2014/10/24. doi: 10.1523/JNEUROSCI.0274-14.2014. PubMed PMID: 25339734.

17. Van Dort CJ, Zachs DP, Kenny JD, Zheng S, Goldblum RR, Gelwan NA, et al. Optogenetic activation of cholinergic neurons in the PPT or LDT induces REM sleep. Proc Natl Acad Sci U S A. 2015;112(2):584–9. Epub 2014/12/31. doi: 10.1073/pnas.1423136112. PubMed PMID: 25548191; PubMed Central PMCID: PMCPMC4299243.

18. Kroeger D, Ferrari LL, Petit G, Mahoney CE, Fuller PM, Arrigoni E, et al. Cholinergic, Glutamatergic, and GABAergic Neurons of the Pedunculopontine Tegmental Nucleus Have Distinct Effects on Sleep/Wake Behavior in Mice. J Neurosci. 2017;37(5):1352–66. Epub 2017/01/01. doi: 10.1523/JNEUROSCI.1405-16.2016. PubMed PMID: 28039375; PubMed Central PMCID: PMCPMC5296799.

19. McCarley RW, Hobson JA. Neuronal excitability modulation over the sleep cycle: a structural and mathematical model. Science. 1975;189(4196):58–60. PubMed PMID: 1135627.

20. Booth V, Diniz Behn CG. Physiologically-based modeling of sleep-wake regulatory networks. Math Biosci. 2014;250:54–68. Epub 2014/02/18. doi: 10.1016/j.mbs.2014.01.012. PubMed PMID: 24530893.

21. Booth V, Xique I, Diniz Behn CG. One-Dimensional Map for the Circadian Modulation of Sleep in a Sleep-Wake Regulatory Network Model for Human Sleep. Siam J Appl Dyn Syst. 2017;16(2):1089–112. doi: 10.1137/16m1071328. PubMed PMID: WOS:000404777600011.

22. Diniz Behn CG, Booth V. Simulating microinjection experiments in a novel model of the rat sleep-wake regulatory network. J Neurophysiol. 2010;103(4):1937–53. Epub 2010/01/29. doi: 10.1152/jn.00795.2009. PubMed PMID: 20107121.

23. Diniz Behn CG, Brown EN, Scammell TE, Kopell NJ. Mathematical model of network dynamics governing mouse sleep-wake behavior. J Neurophysiol. 2007;97(6):3828–40. Epub 2007/04/06. doi: 10.1152/jn.01184.2006. PubMed PMID: 17409167; PubMed Central PMCID: PMCPMC2259448.

24. Robinson PA, Phillips AJ, Fulcher BD, Puckeridge M, Roberts JA. Quantitative modelling of sleep dynamics. Philos Trans A Math Phys Eng Sci. 2011;369(1952):3840–54. Epub 2011/09/07. doi: 10.1098/rsta.2011.0120. PubMed PMID: 21893531.

25. Tamakawa Y, Karashima A, Koyama Y, Katayama N, Nakao M. A quartet neural system model orchestrating sleep and wakefulness mechanisms. J Neurophysiol. 2006;95(4):2055–69. Epub 2005/11/12. doi: 10.1152/jn.00575.2005. PubMed PMID: 16282204.

26. Diniz Behn CG, Booth V. A Fast-Slow Analysis of the Dynamics of REM Sleep. Siam J Appl Dyn Syst. 2012;11(1):212–42. doi: 10.1137/110832823. PubMed PMID: WOS:000302237300008.

27. Diniz Behn CG, Ananthasubramaniam A, Booth V. Contrasting Existence and Robustness of REM/Non-REM Cycling in Physiologically Based Models of REM Sleep Regulatory Networks. Siam J Appl Dyn Syst. 2013;12(1):279–314. doi: 10.1137/120876939. PubMed PMID: WOS:000316865300009.

28. Weber F. Modeling the mammalian sleep cycle. Curr Opin Neurobiol. 2017;46:68–75. Epub 2017/08/26. doi: 10.1016/j.conb.2017.07.009. PubMed PMID: 28841438.

29. Costa MS, Born J, Claussen JC, Martinetz T. Modeling the effect of sleep regulation on a neural mass model. J Comput Neurosci. 2016;41(1):15–28. doi: 10.1007/s10827-016-0602-z. PubMed PMID: 27066796.

30. Schwarz LA, Luo L. Organization of the locus coeruleus-norepinephrine system. Curr Biol. 2015;25(21):R1051–R6. doi: 10.1016/j.cub.2015.09.039. PubMed PMID: 26528750.

31. Luppi PH, Peyron C, Fort P. Not a single but multiple populations of GABAergic neurons control sleep. Sleep Med Rev. 2017;32:85–94. doi: 10.1016/j.smrv.2016.03.002. PubMed PMID: 27083772.

32. Boissard R, Gervasoni D, Schmidt MH, Barbagli B, Fort P, Luppi PH. The rat ponto-medullary network responsible for paradoxical sleep onset and maintenance: a combined microinjection and functional neuroanatomical study. Eur J Neurosci. 2002;16(10):1959–73. PubMed PMID: 12453060.

33. Lu J, Sherman D, Devor M, Saper CB. A putative flip-flop switch for control of REM sleep. Nature. 2006;441(7093):589–94. doi: 10.1038/nature04767. PubMed PMID: 16688184.

34. Weber F, Chung S, Beier KT, Xu M, Luo L, Dan Y. Control of REM sleep by ventral medulla GABAergic neurons. Nature. 2015;526(7573):435–8. doi: 10.1038/nature14979. PubMed PMID: 26444238; PubMed Central PMCID: PMCPMC4852286.

35. Costa MS, Born J, Claussen JC, Martinetz T. Modeling the effect of sleep regulation on a neural mass model. Journal of Computational Neurosciences. 2016;41:15–28.

36. Otchy TM, Wolff SB, Rhee JY, Pehlevan C, Kawai R, Kempf A, et al. Acute off-target effects of neural circuit manipulations. Nature. 2015;528(7582):358–63. doi: 10.1038/nature16442. PubMed PMID: 26649821.

37. Carrera E, Tononi G. Diaschisis: past, present, future. Brain. 2014;137(Pt 9):2408–22. doi: 10.1093/brain/awu101. PubMed PMID: 24871646.

38. Adamantidis AR, Zhang F, Aravanis AM, Deisseroth K, de Lecea L. Neural substrates of awakening probed with optogenetic control of hypocretin neurons. Nature. 2007;450(7168):420–4. doi: 10.1038/nature06310. PubMed PMID: 17943086.

